# Digitization of Electrocardiogram Using Bilateral Filtering

**DOI:** 10.1101/2020.05.22.111724

**Authors:** Hari Sankar S, Anchit Patni, Satish Mulleti, Chandra Sekhar Seelamantula

**Author notes:** Emails: {, }.

## Abstract

Electrocardiogram (ECG) is one of the most basic tools available for the detection of cardiovascular disease (CVD) which has evolved as one of the major causes of death in recent times, spanning countries. Conventionally, ECGs are printed on graph sheets which keep fading away with time. Hence, the preservation of ECG graph sheets is difficult. Digitization of printed ECG graph sheets is a solution to this problem. Digitization also helps in faster analysis and interpretation of ECG signals. Although the previous works have succeeded in smoothing ECG after extracting the signal from graph sheet, the cost amounts to loss of peak amplitude characteristics leading to wrong diagnosis. In this work, we propose a digitization technique to extract ECG signals from graph sheets. The proposed bilateral filter based method achieves a high degree of smoothness with all the peak characteristics intact when compared to other implementations using Butterworth filters. With this technique, it is now possible to create extensive databases similar to MIT–BIH from printed ECG graph sheets. A high level of automation is achieved, and results verified with over 60 ECGs and compared.

## 1. INTRODUCTION

Electrocardiography is the process of recording the electrical activity of the heart using electrodes placed on the chest, arms, and the legs of a subject. In this non-invasive medical procedure, the action potentials of the excitable cardiac cells are plotted on a voltage versus time graph called electrocardiogram (ECG). The ECG waveform is characterized by a series of waves whose morphology and timing information are used for measuring heart rate [1], presence of worn out heart muscle cells, effect of drugs [2], monitoring implanted pacemakers [3], diagnosing cardiovascular diseases like arrythmia [4], coronary events like myocardial infarction [5], etc.

Conventionally, the ECG is printed on a thermal paper for detailed examination by medical practitioners. This leads to a large volume of ECG sheets which in turn results in a tedious and error prone process of examining and retrieval of ECG. Further, sharing this bulky collection of sheets among doctors is a difficult task. Digitization of printed ECG graphs will help in efficient storing and sharing of recorded ECG. It also aids in automatic feature extraction from ECG and diagnosis.

It is extremely difficult to find reliable databases of ECG like the MIT-BIH arrythmia database in India. This database is based on digitization methods available at that time [6] and is prone to errors due to a lot of manual work [7]. Hence, digitization is important for both clinical and research purposes. In this paper, we propose a novel ECG digitization methodology. Before going into the details of the proposed method we briefly review the related literature.

### 1.1. Literature Survey

After the MIT-BIH arrhythmia database was created, a lot of research work in digitization of ECG paper recordings has been done [8]. Preserving the smoothness of the digitized ECG and the peaks of the PQRS and T waves remains a difficult task. Morales et al. [9] have managed to produce an improvement by reducing the magnitude of the overall rootmean-square error, but the error still remains significant. In general, the first step in digitization is the separation of the ECG waveform from the grid lines using image intensity histogram [10]. This step is prone to errors as it is completely depended upon the user’s ability to observe and select the threshold. Syeda et al. [11], Wang et al. [12] and Kao et al. [13] proposed the use of morphological operators which adds its own artifacts to the original waveform.

Related works on converting ECG signal from graph sheet to digital format focus only on particular aspects of ECG, have significant loss of data in the waveform peaks and also are specific to a particular ECG paper recording. Filtering techniques like Laplacian filtering [14] which is applied for reducing the background noise, causes distortion in the ECG signal. Waveform contour detection by gradient [15] is another approach that depends heavily on true detection of each and every contour. Widman et al. [16] converts the graph sheet to binary in the very first stage itself which is not encouraged when accuracy is a constraint. Leung and Zhang [17] have followed a delta-sigma modulation technique but the reconstructed signal is again passed through a Butterworth filter which affects the peaks and valleys. As we can see, in the previous works, there is a trade-off between smoothness and the amplitude of the peaks, so we are proposing a bilateral filter based method to achieve a high degree of smoothness along with all the peaks preserved.

### 1.2. This Paper

ECG corrupted with noise can lead to wrong diagnosis. Data loss at the waveform peaks still remains a common problem [8]. The motivation for this research is to adopt a novel method to obtain a smooth and digitized ECG waveform from the paper recordings with the peaks preserved. Bilateral filter has evolved as a very effective edge preserving smoothing filter for image processing applications [18] [19] but its potential remains untapped in the case of one-dimensional signals. Bilateral filter seems to be an ideal candidate to serve the purpose. The proposed method is automated and does not require any human intervention. This method has been tried on more than 60 real ECG paper recordings and the digitized waveforms compared with the original scanned ECG recordings for validation.

### 1.3. Paper Organization

The organization of paper is as follows: In section 2, we briefly discuss the bilateral filter. In section 3, the proposed ECG digitization technique is discussed. In Section 4, we note the results obtained and conclusions are discussed in section 5.

## 2. MATHEMATICAL PRELIMINARIES

Bilateral filter is a non-linear edge preserving filter often used in image denoising. The filter consist of a domain kernel and a range kernel. The weights in the domain kernel depends on the distance between the present sample and its neighboring samples, whereas the weights in the range kernel are dependent on the difference between the amplitudes of the present sample and the neighboring samples. Suppose we are given a sequence of measurements *y* (*n*) = *x*(*n*) + *w*(*n*), *n* ∈ ℤ, where *x*(*n*) is the desired clean signal and *w*(*n*) is noise term. By applying the bilateral filter, the estimation of *x*(*n*) from *y* (*n*) is given by

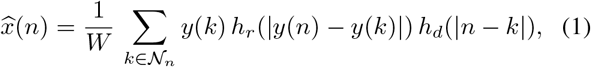

where 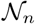 is neighborhood around the sample location *n* and

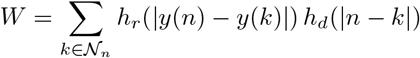

is normalization factor. In (1), *h_r_*(·) and *h_d_*(·) are range and domain kernels, respectively.

## 3. PROPOSED METHOD

The method proposed involves the extraction of ECG waveform from the paper recordings, digitizing itandthensmoothing of the obtained waveform along with preservation of peaks. The steps of the algorithm are explained as follows:

### 3.1. Scanning a paper ECG

At first, the available ECG graph sheet is scanned at a high resolution of 600 dpi and saved in TIF file format, which retains important features of the signal. Fig. 1 shows one such scan. In the succeeding steps, we show the results on this particular scan.

**Fig. 1.**
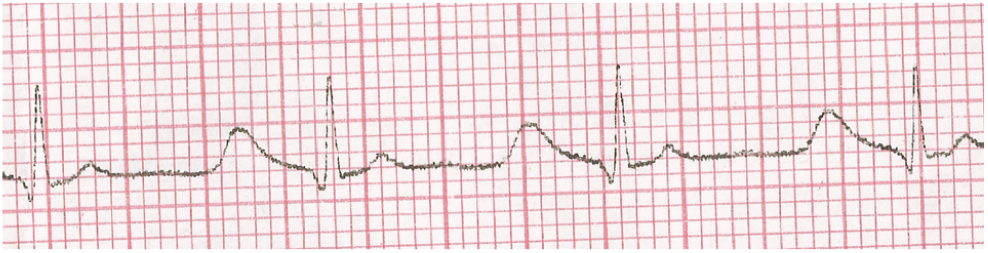
Scanned paper ECG

### 3.2. De-skewing and thresolding

As scanning a graph sheet is prone to human errors in terms of skew angle, we need to de-skew the image before processing it further. This is done with the help of Hough transform [20]. The phase of the Hough peak gives the angle for the re-orientation (rotation). De-skewed image is as shown in Fig. 2.

**Fig. 2.**
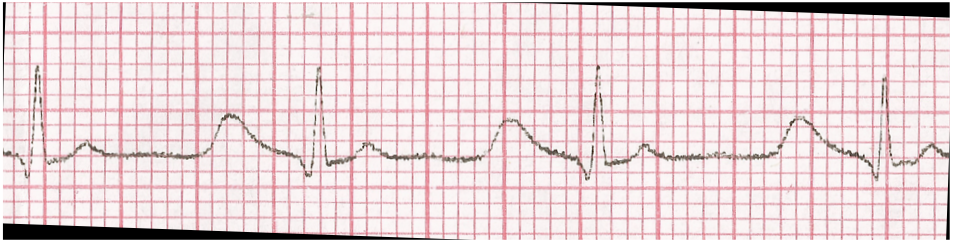
Deskewed image of the image shown in Fig. 1.

Instead of using the threshold based on the color intensity histogram, we have applied Otsu method to find the threshold to separate the grid lines from the data waveform followed. The resulting image contains isolated dark pixels which are annihilated by applying median filter. The resulting waveform free from the grid lines is shown in Fig. 3.

**Fig. 3.**
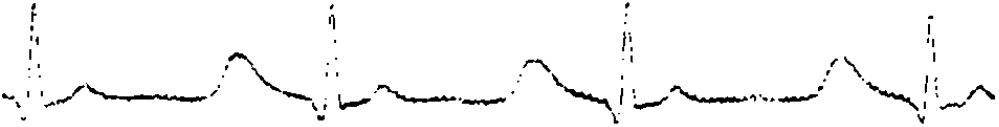
Extracted waveform from Fig. 2 by applying Otsu thresholding and median filtering.

### 3.3. Digitization

The waveform extracted in the previous step is in the form of a two-dimensional matrix. The next step is to extract the waveform from this matrix or image and convert it to a onedimensional signal. After extracting the waveform

For digitization in our algorithm, all the dark pixels along the column as in Fig. 3 are selected and the median of the distances from the bottom of the image of all the dark pixels are stored as a vector. Plotting this followed by interpolation will provide us the digitized ECG signal which does not appear very smooth.

### 3.4. Smoothing and peak preservation

In order to achieve a high degree of smoothness in ECG and to preserve all the peak amplitudes to a very high extent, we use bilateral filtering. In bilateral filtering, the filtered amplitude depends on a weighted combination of neighboring amplitudes. In this work, we have used a Gaussian function as range and domain kernels with zero mean and standard deviations *σ_r_* and *σ_d_*, respectively. We chose 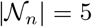.

Performance of bilateral filtering depends mainly on two parameters: firstly the domain kernel parameter *σ_d_*, and secondly, the range kernel parameter *σ_r_*. As *σ_r_* increases, the filter gradually approaches a normal Gaussian filter because the range kernel widens and flattens. As either of the parameters approaches zero, no smoothing occurs. As a consequence, increasing *σ_d_* will not smoothen the peaks as long as *σ_r_* is less than the amplitude of the peak to be preserved. Increasing *σ_d_* has limited effect unless *σ_r_* is also increased.

By applying the filter for different combinations of *σ_r_* and *σ_d_* on 60 ECG recordings, we experimentally found that *σ_d_* should lie in between one and two. While fixing *σ_r_*, it is important that *σ_r_* should be less than the difference in amplitudes of any two adjacent points at the peak. Satisfying these conditions for obtaining the desired output, value of *σ_r_* turns out to be less than or equal to 5. Experimentally, we found that for these ranges of *σ_r_* and *σ_d_* we have the desired peak amplitude preservation and smoothness.

The proposed ECG digitization software and MATLAB code for the same are available for reference and can be downloaded from www.ee.iisc.ernet.in/downloads/software.

## 4. RESULT

Fig. 5 shows the bilateral filtered output of the ECG paper shown in the fig. 1. Fig. 6 shows the superimposition of the bilateral filtered output and scanned ECG graph paper. Fig. 7 shows the zoomed portion of the superimposition for a better view. Fig. 8 and fig. 9 show the comparison between the Butterworth filter and bilateral filter. It was upheld that the Butterworth filter gives a better denoising of ECG as compared to other filters [17]. But Fig. 8 and 9 shows clearly that our bilateral filter based approach gives a more accurate representation of digitized ECG along with smoothing it. Lower order Butterworth filters greatly deviate from the ideal brick wall response and hence, poses a high risk of hiding the elevated ST–segment indicating anteroseptal myocardial infarction [21] or can even give a false alarm of a pathologic Q– wave indicating a previous myocardial infarction [22]. Bilateral filter comes to our rescue here. Fig. 9 establishes that the peaks are preserved to a great extent by the bilateral filtering while denoising ECG; peak degradation by the Butterworth filter is very high.

**Fig. 4.**
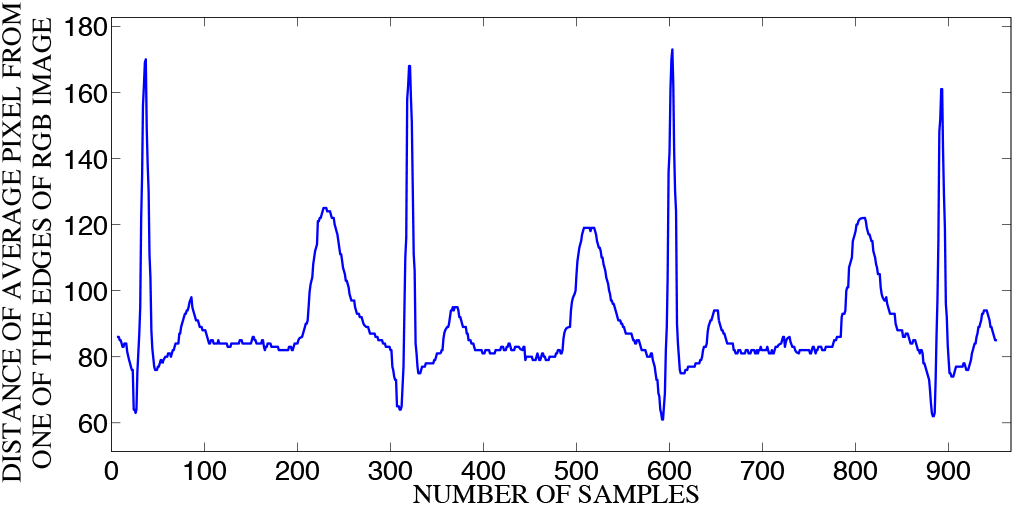
Digitized ECG waveform

**Fig. 5.**
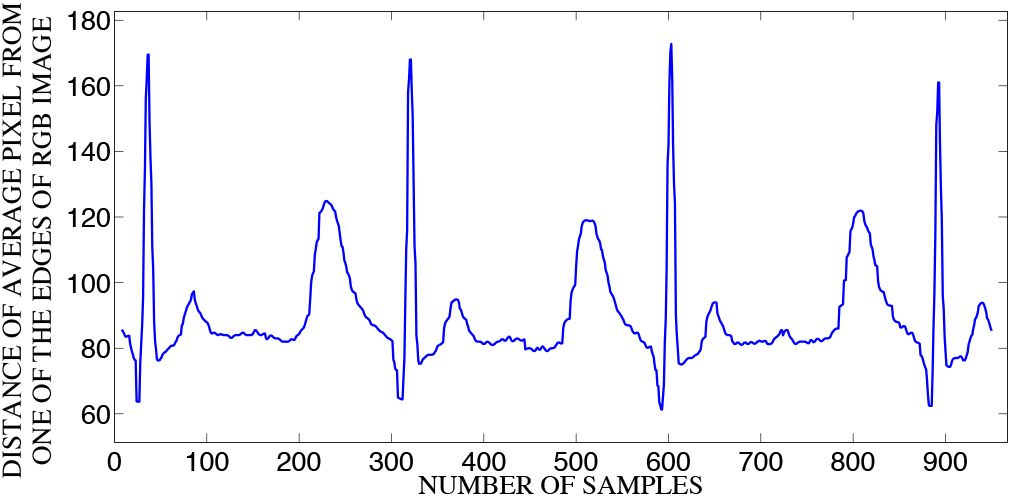
Bilateral peak preserved smoothened output

**Fig. 6.**
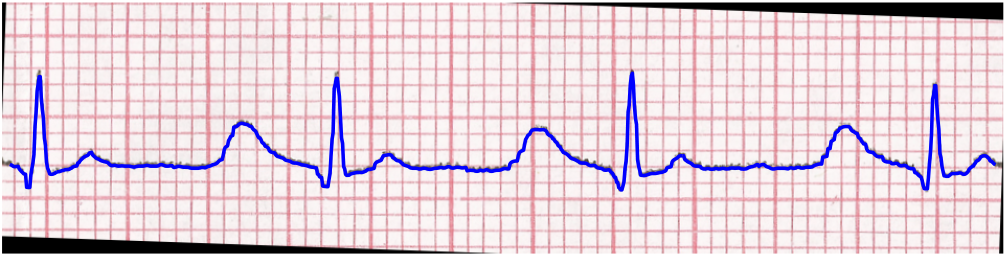
Superimposition of the bilateral filtered output and scanned ECG graph paper

**Fig. 7.**
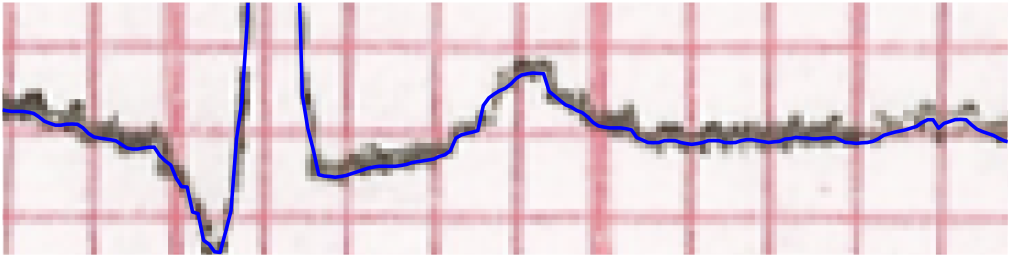
Zoomed view Fig: 6

**Fig. 8.**
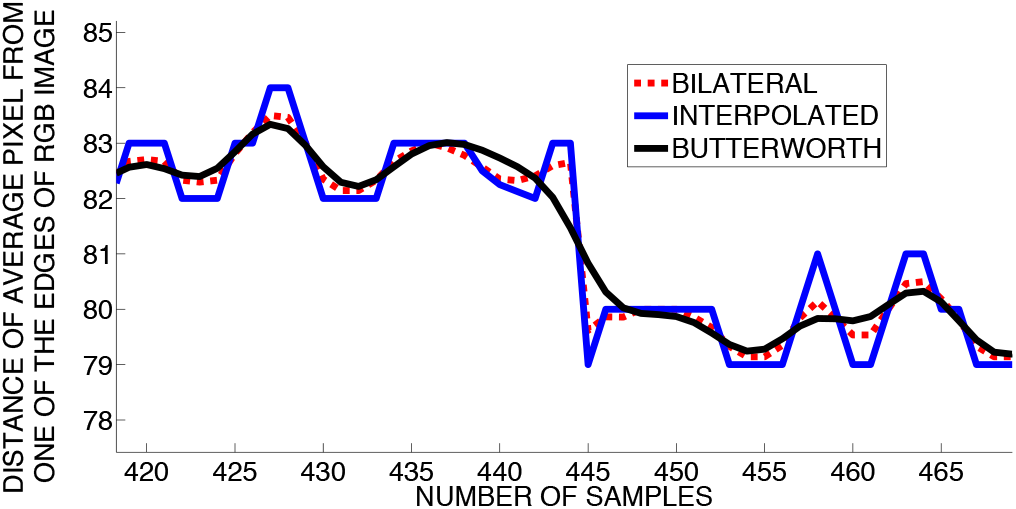
Shows the comparison of smoothness of Butterworth and bilateral filtered outputs

**Fig. 9.**
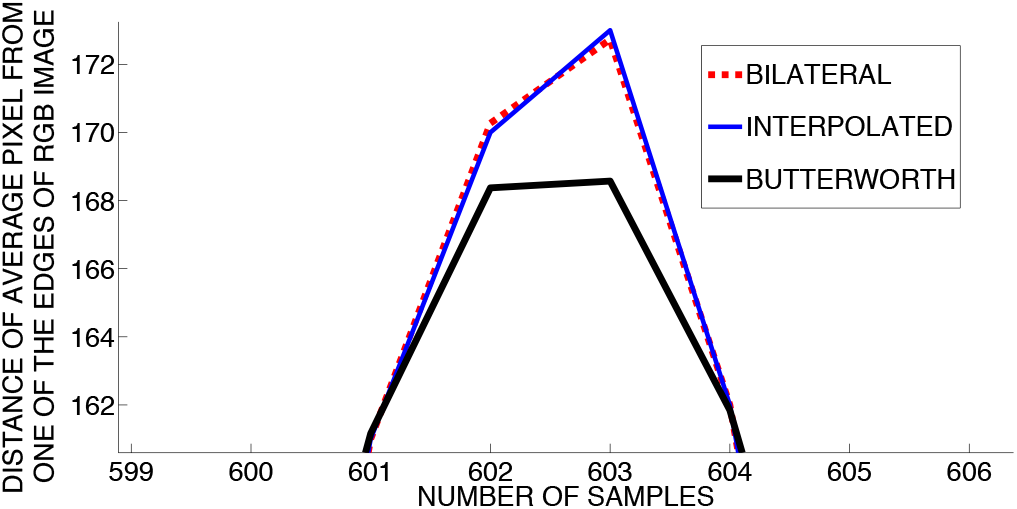
Shows the comparison between Butterworth and bilateral on peak preservation

## 5. CONCLUSIONS

The bilateral filter provides an efficient and reliant tool for ECG smoothing without affecting the peak amplitudes of the waveform. The Butterworth filter, comparatively unreliable when it comes to the combo of relative smoothness and peak preservation can henceforth be replaced with the bilateral filter. The desired amount of smoothness can be varied by changing σr and σd within the limits. The digitization algorithm is complete in itself and has been validated with 60 ECGs. The Otsu thresholding makes this adaptive to a wide range of image quality.

## Notes

### Competing Interest Statement

The authors have declared no competing interest.

